# When neighbours play a role: a systems-level analysis of protein interactions conditioning cancer driver mutation effects

**DOI:** 10.1101/2025.01.18.633728

**Authors:** João AI Miranda, Márcia FS Vital, Margarida Carrolo, António Quintela, Francisco R Pinto

## Abstract

**Background:** Cancer is driven by the accumulation of somatic mutations, including driver mutations that confer a selective advantage to cancer cells. Driver proteins operate within complex interaction networks, and their activity is conditioned by neighbour proteins. Understanding the interplay between driver mutations and the expression of their neighbour proteins can provide insights into cancer biology and potential therapeutic targets.

**Methods:** We assessed associations between expression of neighbour proteins and driver mutation status, comparing both between and within cancer types. We further evaluated if neighbours were enriched in significant associations with multiple drivers and characterised the impact of neighbour expression on overall survival for all cancer types.

**Results:** We found a significant correlation between the number of driver associations a neighbour gene has and the number of sign-coherent survival associations, particularly for neighbours enriched in positive associations, where high neighbour expression correlated with increased driver mutations and poorer survival. We identified 119 neighbours enriched in positive driver associations with at least two unfavourable survival associations and 25 neighbours enriched in negative driver associations with at least two favourable survival associations.

**Conclusions:** Our study systematically identified neighbours associated with driver mutation status. Complementary evidence from survival analysis and the literature suggests that neighbours enriched in driver associations can be further explored as drug target candidates.

**Significance Statement:** Cancers are caused by mutations in driver genes. The impact of those mutations in the cell can be influenced by other proteins in the cell that physically interact with the mutated protein. In this work, we analysed cancer patient data to uncover such neighbour proteins that may influence the outcome of driver mutations. We discovered a subset of neighbour proteins that are associated with mutations in multiple cancer drivers and, simultaneously, are associated with changes in survival times of patients for multiple cancer types. These neighbour proteins may help explain why some driver mutations are more common in certain cancer types. We also propose that these neighbour proteins should be explored as candidate drug targets for cancer therapy.

## Introduction

Over the last decades, our understanding of oncogenesis has significantly expanded. The discovery of cancer driver genes in the 1970s has contributed to the advancement of targeted anticancer therapies and, more broadly, to the identification of genomic biomarkers for predicting prognosis and treatment response (1). Likewise, the arrival of genomic sequencing of tumours has led to a remarkable collection of disease-driving mutations, offering valuable insights into cancer genomics (2).

Although 91% of tumours carry at least one gene driver mutation (3), these mutations alone are insufficient to fully explain the process of cell transformation into tumour cells (4). Furthermore, research in healthy individuals has shown that some cancer driver mutations are already present in non-transformed cells across different somatic tissues, supporting the belief that mutations in cancer driver genes *per se* are not enough to promote tumorigenesis (5, 6).

The cellular context in which a driver mutation occurs, namely the physical interactors that are present, can condition the impact of the mutation on the cell phenotype (7). Protein-protein interactions (PPIs) play a crucial role in both physiological and pathological processes, such as cell proliferation, signal transduction and apoptosis (8, 9). Consequently, mapping of cancer-related PPIs is an ongoing effort to better understand disease and contribute to the development of precision medicine approaches (10–14).

In the last decades, PPIs have gained growing attention and emerged as therapeutic targets for both haematological and solid tumours (15, 16). Examples of therapeutic approaches targeting cancer driver PPIs include inhibitors of MDM2/p53, KRAS/PDEδ and c-Myc/Max interactions (15).

Despite these advances, the systematic investigation of the relationship between driver mutations, neighbour gene expression, and patient survival remains largely unexplored. This study aims to address this gap by analysing the association between driver mutations and the expression of their neighbour genes in a pan-cancer cohort. We hypothesize that the expression of neighbour genes can modulate the phenotypic effects of driver mutations and influence patient survival. If, for example, the driver protein is a protein kinase, the effects of the mutation will be different depending on the set of neighbour phosphorylation targets that are expressed in the cell type where the mutation occurred. Alternatively, neighbours can influence the stability of the driver, and consequently its steady-state abundance, as is the case of the PPIA stabilising effect on the oncogene NRF2 (17). Moreover, we hypothesize that when such neighbour effects exist, we can identify them through correlations between neighbour expression and driver mutation status in cohorts of cancer patients. By integrating genomic, transcriptomic and interactomic data, we aim to identify novel therapeutic targets and gain a deeper understanding of the complex interplay between driver mutations and their protein interaction context in cancer.

## Results

### Rationale and computational workflow

In this study, we sought to find physical interactors (hereafter called “neighbours”) of cancer driver proteins that influence how driver mutations cause cancer. We reasoned that if a neighbour affects the phenotypic consequences of driver mutations in a cancer cell, it should be possible to identify a correlation between the neighbour gene expression and the presence of driver mutations in cancer patient data. Pharmacologically targeting these neighbours might lead to an attenuation or even reversion of the driver mutation effects. With this in mind, we designed a computational workflow (Figure 1) to analyse cancer patient genomic and transcriptomic data (from the TCGA Pan-Cancer cohort (3, 18) and evaluate Cancer Driver PPIs (known cancer driver genes extracted from NCG (19) and PPIs from Biogrid (20), String (21), HuRI (22), Omnipath (23) and APID (24), global network properties available in supplementary table S1).

**Figure 1.**
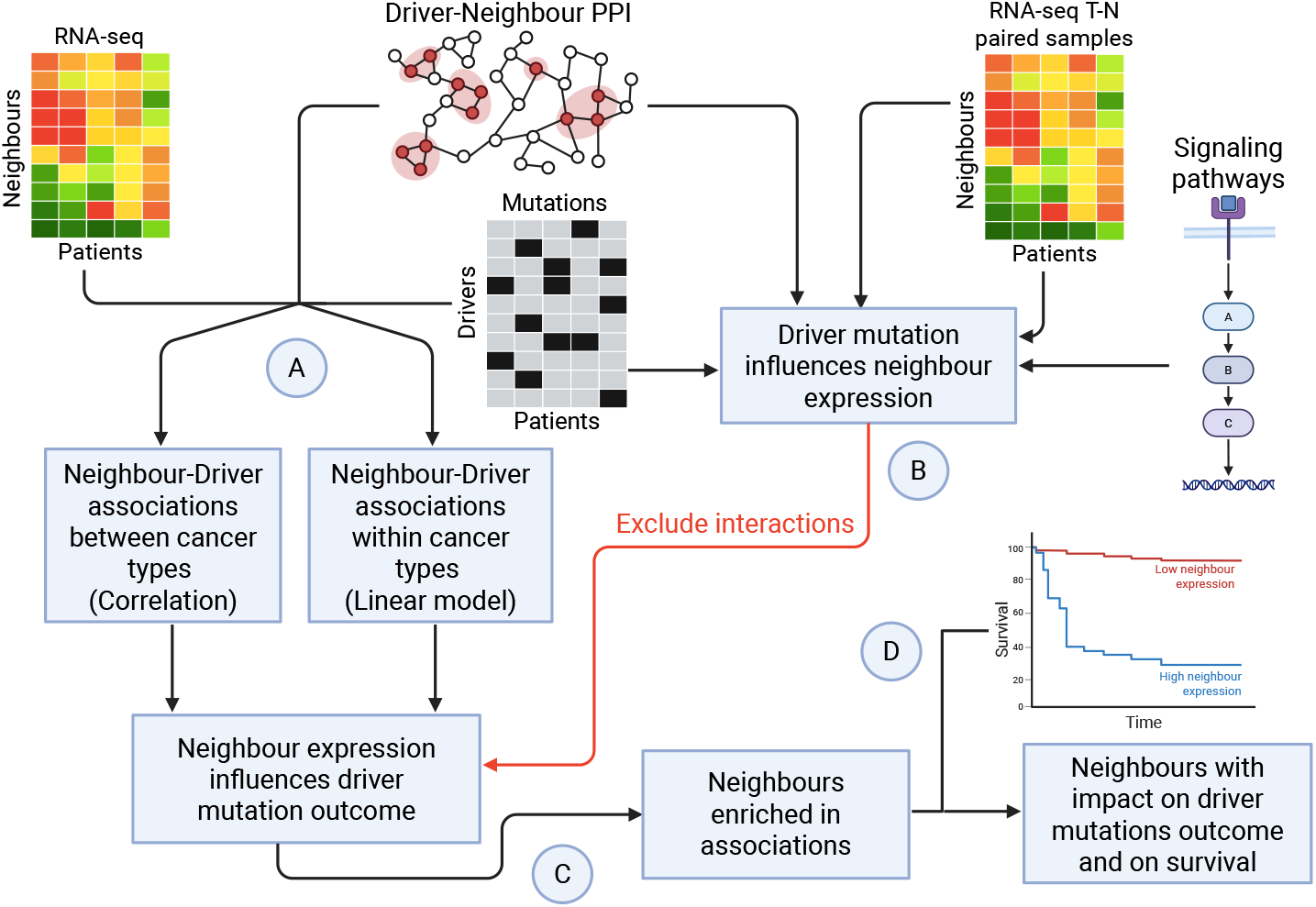
Computational analysis workflow. (A) Using cancer patient mutation data, PPI between cancer drivers and neighbour proteins, and gene expression data from the same patients, we evaluated two types of statistical association between neighbour expression and driver mutation status, one comparing between cancer types and the other comparing within cancer types. (B) To avoid statistical associations where change in gene expression is caused by driver mutations, we also evaluated associations between the difference in neighbour expression between tumour and normal matching tissues and driver mutation status. Statistically significant associations found with paired tumour-normal samples were excluded from further analyses. Additionally, we searched signalling and transcriptional regulatory networks for directed pathways linking the driver and the neighbour genes. If those pathways had a length of 2 or fewer interactions, the driver-neighbour pair was also excluded from further analyses. (C) Next, we selected neighbours that had more driver associations than expected by chance, using analyses with randomly permuted expression datasets to estimate the random expectation. (D) Finally, we checked a previously available survival analysis study for neighbours with a significant impact on patient overall survival for several cancer types. These results allowed us to select neighbours that were simultaneously enriched in driver association and had survival associations with coherent effect signs (positive or negative association with driver mutations concurrently with a decrease or an increase in overall survival times, respectively). Created in BioRender. Gama, M. (2025) https://BioRender.com/o45×904.

The expression of neighbours can vary between tissues and between individuals. Therefore, we followed two approaches to detect neighbours associated with driver mutation status: one comparing neighbour expression between cancer types (BCT) and the other comparing neighbour expression within cancer types (WCT) (Figure 1 A). Both methods considered the possible effect of the tumour mutational burden (TMB) in each individual or cancer type. But the detection of these associations does not imply a causal relationship. Three types of causal relationships can originate these associations: 1) the driver mutation induces changes in neighbour expression; 2) the neighbour expression influences the phenotypic outcome of the driver mutation; or 3) a third variable influences simultaneously the occurrence of mutations in the driver and the neighbour expression. We aim to find candidate neighbours with significant associations with cancer drivers due to the second type of causal relationship. The third type of causal relationship can only be excluded with experimental tests, but the first type can be excluded with 1) available cancer patient data and 2) known signalling pathways and transcriptional regulations (Figure 1 B). First, using a cohort of patients in which gene expression was measured in both tumour and adjacent normal tissue, we could infer the effect of the driver mutation by quantifying the difference in the neighbour expression between tumour and normal tissues (tumour/normal tissue comparison). If the difference is the same in patients with and without driver mutations, the mutation does not have a significant impact on the neighbour expression. Additionally, we identified short directed pathways connecting driver to neighbour genes. These directed pathways were extracted from known signalling networks (available in Omnipath (23)). The final step of each directed pathway was always a transcriptional regulatory interaction targeting the neighbour gene (extracted from the DoRothEA resource (25)). The existence of these short, directed pathways suggests that the driver can influence the neighbour expression. Indeed, we determined that driver neighbour pairs connected by directed pathways of length 1 or 2 were significantly more likely to exhibit a significant difference in the tumour/normal tissue comparison (Supplementary table S2). Neighbour-driver PPIs that 1) were not significant in the tumour/normal tissue comparison and 2) were not connected by directed pathways of length 1 or 2, were kept for further analysis. Due to the very large number of neighbour-driver PPIs under evaluation (401402 in the WCT analysis and 320055 in the BCT analysis) it is very difficult to apply a statistical cut-off that ensures a low fraction of false positives without drastically losing statistical power (meaning that many false negatives are produced). Therefore, we focused on identifying neighbours that were involved in a larger number of significant associations with drivers than expected by chance (Figure 1 C). Lastly, we compared our results with a previously available survival analysis (26, 27) to determine whether the identified neighbours might have an impact on disease progression and severity (Figure 1 D).

### Neighbour-Driver associations are detectable between and within cancer types

We detected BCT associations by computing the Spearman rank correlation coefficient between the fraction of individuals with mutations in a driver (normalised by the average TMB of each cancer type) and the mean expression of the driver neighbour for each cancer type. Figure 2A shows one example of a positive BCT association between neighbour expression and driver mutation. This correlation and the associated statistical test were computed for all driver-neighbour PPIs. To detect WCT associations, we applied linear models to predict neighbour expression using driver mutation status, TMB and cancer types as predictor variables. Figure 2B shows one example of a positive WCT association between neighbour expression and driver mutation. When model coefficients for driver mutation status are significantly different from zero, we can infer that neighbour expression is consistently distinct between individuals with and without driver mutation for multiple cancer types. The obtained distribution of p-values in both analyses showed an enrichment for lower p-values, contrasting with the uniform distribution obtained when the same tests were applied with permuted expression datasets (Figure 2C and 2D). This departure from the uniform distribution confirms the existence of driver-neighbour associations between neighbour expression and driver mutation status, both across and within cancer types. Considering significant associations with a p<0.05, we obtained 30762 BCT and 30361 WCT associations.

**Figure 2.**
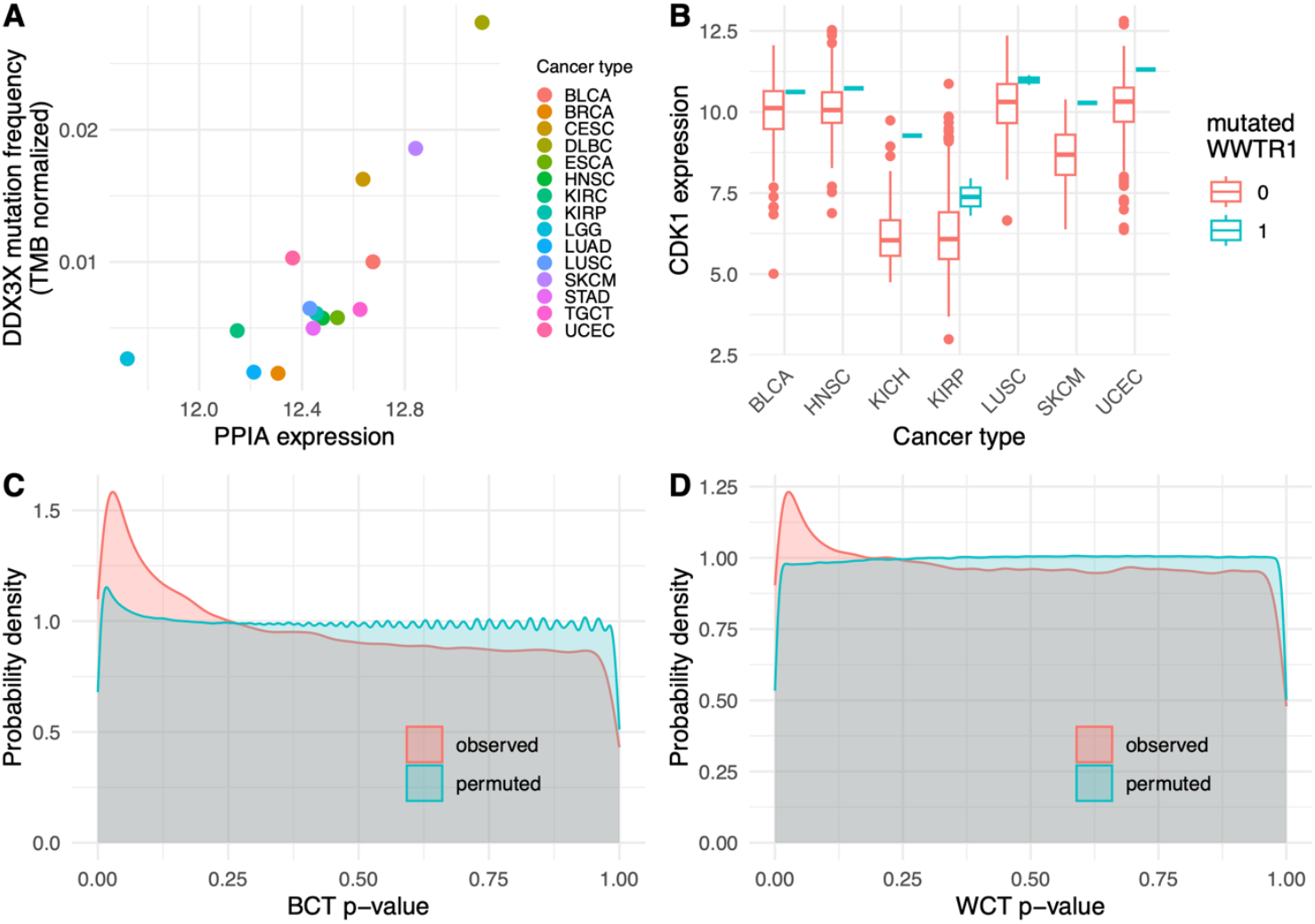
Driver-neighbour associations. (A) Example of a significant BCT statistical association between neighbour (*PPIA* in the example) expression and the frequency of driver mutation (*DDX3X* in the example). Each dot represents data from a different cancer type. (B) Example of a significant WCT statistical association between neighbour (*CDK1* in the example) expression and driver (*WWTR1* in the example) mutation status. Neighbour expression values are represented as box plots according to cancer type and driver mutation status. In (A) and (B) expression values are log2(normalised TPM + 1). All significant BCT and WCT associations can be explored at https://joaoaimiranda.shinyapps.io/driver_neighbours_dashboard/, including correlation and regression coefficients and p-values. (C) Probability density distributions of Spearman correlation test p-values obtained using observed data or randomly permuted expression datasets. (D) Probability density distributions of mutation status linear model coefficient p-values obtained using observed data or randomly permuted expression datasets

Although both analyses took into account TMB, results could be biased by complex differences in mutational signatures across subgroups of samples. To determine whether our findings were being influenced by such effects, we employed the Coselens method (28) to analyse 10 WCT associations (chosen from the top-ranked associations according to statistical significance, avoiding associations with repeated drivers or neighbours). This method compares the mutation frequency of a gene across two sample groups, taking into account a fitted model of the mutational profiles in each sample group. In our case, the two sample groups were defined by their level of neighbour gene expression. For all 10 associations, we were able to detect evidence of significant conditional selection of driver mutations by neighbour expression in at least one cancer type (Supplementary data file 1). Moreover, the promotion or inhibition of driver mutations was concordant with the sign of the WCT associations.

On the other hand, BCT associations could be due to a match between the neighbour expression tissue specificity and the cancer types with higher (or lower) driver mutation frequency. In that scenario, the neighbour could be associated with the driver mutation frequency, but not necessarily involved in the tissue specific factors that condition the effects of driver mutations. These tissue specificity induced BCT associations would be due to a small number of cancer types with high (or low) driver mutation frequency and very high (or very low) neighbour expression. To rule out the possibility that these phenomena were prevalent in our dataset, we partitioned the cancer types for each driver-neighbour pair into two subsets based on driver mutation frequency, and calculated the BCT correlations for each subset separately. For pairs with a significant BCT association, the signs of the low-mutation (or high-mutation) BCT correlation were in agreement with the correlation using all cancer types in more than 80% of the associations (Supplementary table S3). This sign concordance was lower for non-significant associations (around 60%). This suggests that significant BCT associations are supported by robust trends that are not only due to a few cancer types that happen to coincide with the tissue specific expression of neighbour genes. Additionally, we checked the tissue specificity of neighbour genes (Supplementary figure S1) and found that neighbours enriched in BCT associations tend to have a lower tissue specificity index, meaning that they are more broadly expressed across numerous tissues. This observation further supports the notion that our of significant BCT associations are not due to a spurious neighbour tissue specificity effect.

The list of driver-neighbour PPIs evaluated included 13748 (for BCT) and 14236 (for WCT) neighbours, but only around 55% of them were involved in significant BCT associations and 60% in WCT associations (Table 1). We compared these percentages with what would be expected by chance if the same number of significant associations were attributed randomly among all driver-neighbour PPIs. While the number of neighbours with significant WCT associations was close to what would be expected by chance, the number with significant BCT associations was significantly lower than expected. Regarding driver participation in significant associations, the fraction was higher than for neighbours (above 80% for both BCT and WCT) but significantly lower than expected by chance for both types of association (Table 2). These deviations from randomness suggest that some drivers (and some neighbours in the case of BCT associations) are more likely to be involved in significant driver-neighbour associations.

**Table 1.**
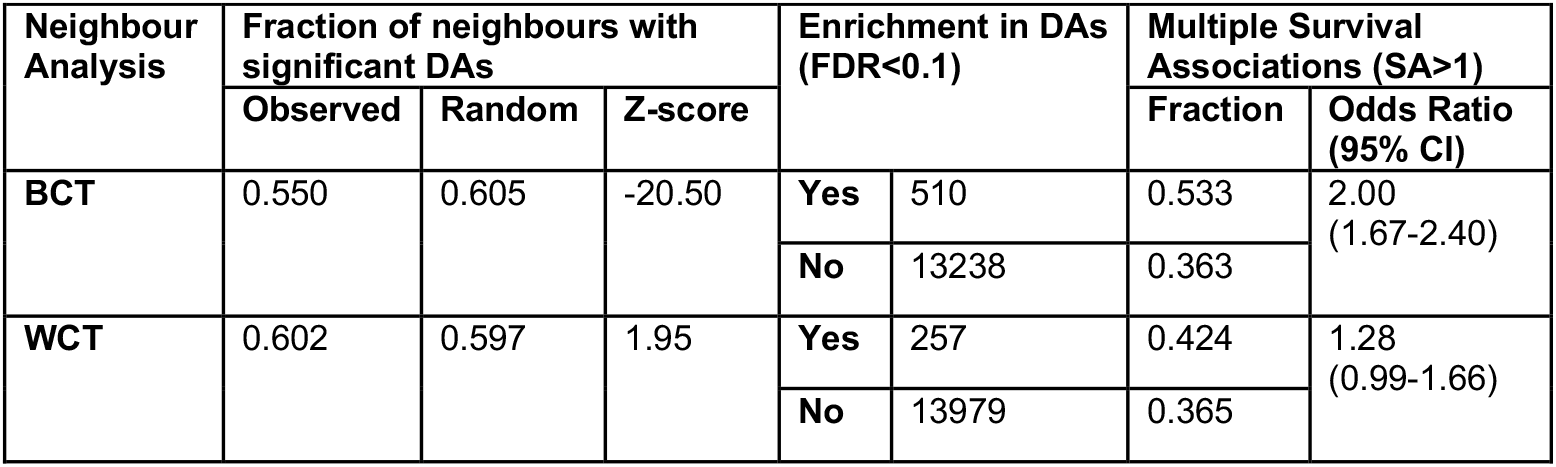
Characterisation of neighbours with significant driver associations.

**Table 2.** Characterisation of drivers with significant neighbour associations.

| Driver Analysis | Fraction of drivers with significant NAs |  |  | Enrichment in NAs (FDR<0.1) |  | Canonical drivers |  |
| --- | --- | --- | --- | --- | --- | --- | --- |
|  | Observed | Random | Z-score |  |  | Fraction | Odds Ratio (95% CI) |
| **BCT** | 0.840 | 0.899 | -13.90 | **Yes** | 403 | 0.241 | 1.43 (1.10-1.86) |
|  |  |  |  | **No** | 2000 | 0.182 |  |
| **WCT** | 0.821 | 0.879 | -13.78 | **Yes** | 42 | 0.810 | 19.50 (8.80-49.09) |
|  |  |  |  | **No** | 3005 | 0.179 |  |

We then evaluated if some neighbours or drivers were involved in more associations than expected by chance. For that, we repeated all analyses with 100 datasets where expression profiles were randomly permuted between all patients. In this way, the eventual associations between neighbour expression and driver mutation status were abolished, and the resulting distribution of the number of associations per neighbour (or driver) was used to evaluate a null model random expectation. For each neighbour and driver, the number of significant associations was transformed into a z-score. Similar z-scores were computed for the permuted datasets, allowing us to set a z-score threshold above which a false discovery rate lower than 0.1 was ensured. This resulted in the identification of 510 and 257 neighbours enriched in BCT and WCT driver associations (DA), respectively (Table 1). For drivers, 403 and 42 were enriched in BCT and WCT neighbour associations (NA), respectively (Table 2). Among the cancer drivers included in our study, 591 are considered well known canonical drivers, while the remaining 2456 drivers have been identified in genomic studies of cancer patient cohorts (collected and annotated in the NCG resource (19)) but the experimental evidence of their driver role is less well established. The subsets of drivers enriched in BCT and WCT NA have significantly higher proportions of canonical drivers than expected (Table 2). This suggests that these driver-neighbour associations are characteristic of true cancer drivers. Some of the canonical drivers are labelled as oncogenes or tumour suppressor genes. Oncogenes become overly active due to driver mutations, while mutations in tumour suppressor genes result in the loss of their function. For drivers enriched in WCT NA, both oncogenes and tumour suppressor genes tend to have higher z-scores than drivers with unknown role, while for BCT NA, only tumour suppressor genes tend to have higher z-scores (Supplementary figure S2).

### Neighbours enriched in BCT associations tend to have positive sign-coherent effects

We classified enriched neighbours as “positive” (when at least 80% of their driver associations are positive), “negative” (when at least 80% of their driver associations are negative) or “mixed” (remaining cases). A higher expression of positive neighbours is associated with the tumorigenic effect of driver mutations. So, we can infer that these neighbours may have a cellular function that is intrinsically oncogenic. Conversely, negative neighbours should have tumour suppressor roles. Mixed neighbours have a context-dependent role according to which drivers are mutated. Neighbours enriched in BCT DA show a different pattern of association signs when compared with neighbours enriched in WCT DA (Figures 3A and 3B). A high proportion of neighbours enriched in BCT DA had the majority of their associations with a coherent sign, either positive or negative, while neighbours enriched in WCT DA tend to have a mixture of positive and negative associations.

**Figure 3.**
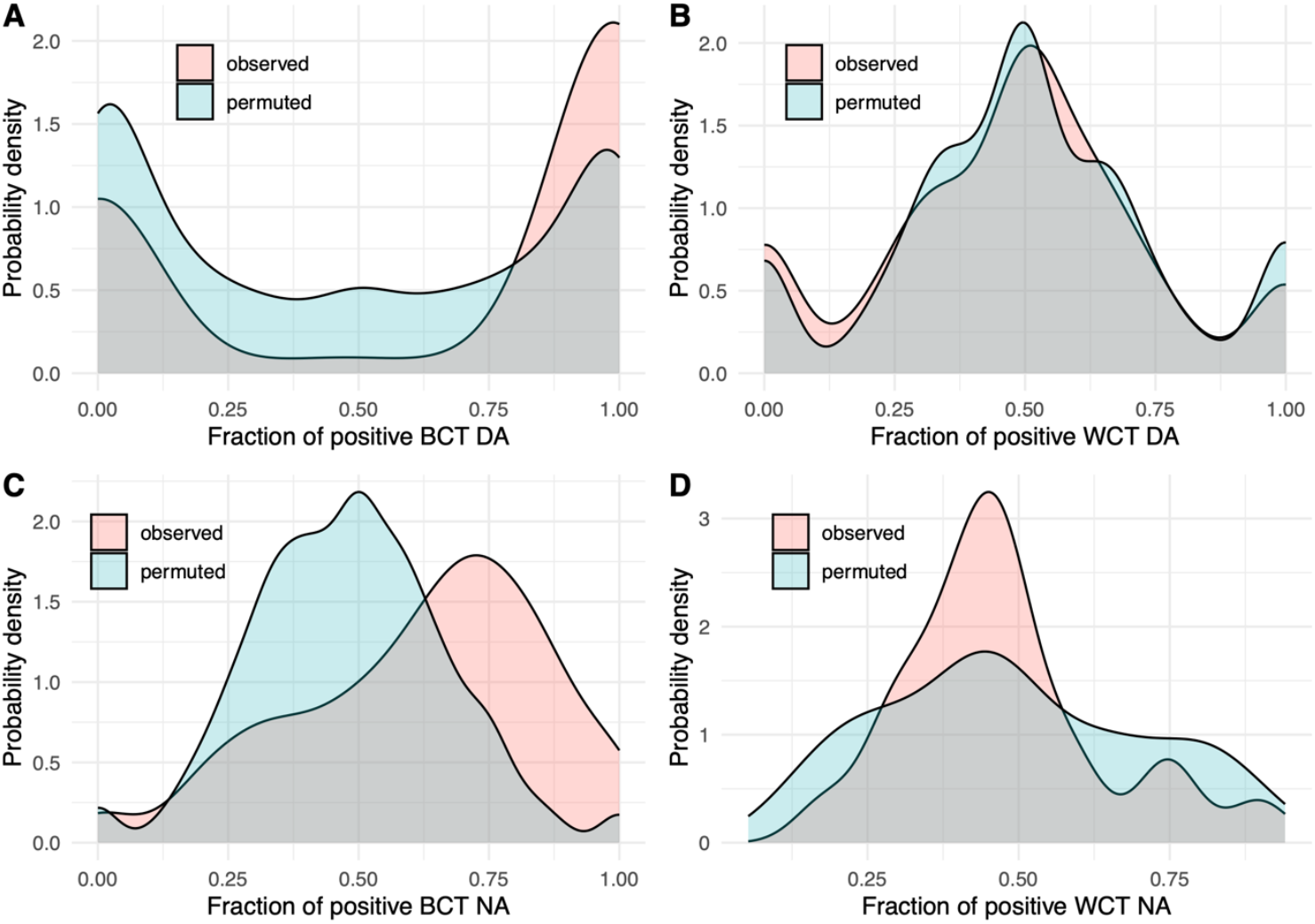
Probability density distributions of the fraction of positive associations. (A) Probability density distributions of the fraction of positive associations for neighbours enriched in between cancer type (BCT) driver associations (DA) and comparison with the corresponding distribution in analyses using permuted expression datasets. (B) Probability density distributions of the fraction of positive associations for neighbours enriched in within cancer type (WCT) DA and comparison with the corresponding distribution in analyses using permuted expression datasets. (C) Probability density distributions of the fraction of positive associations for drivers enriched in BCT neighbour associations (NA) and comparison with the corresponding distribution in analyses using permuted expression datasets. (D) Probability density distributions of the fraction of positive associations for drivers enriched in WCT DA and comparison with the corresponding distribution in analyses using permuted expression datasets. All distributions obtained with permuted datasets were matched with observed distributions by the number of significant associations per neighbour or driver.

Part of this difference in association sign pattern is probably due to differences in the structure of correlations between driver mutation patterns. Two drivers that tend to be simultaneously mutated in the same individuals will also be positively correlated in their cancer type mutation frequency profile. On the other hand, drivers with a positive correlation of cancer type mutation frequency are not necessarily positively correlated at the individual level. This is expected for drivers that are part of the same pathway and have a mutual-exclusivity mutation pattern, which should be positively correlated at the cancer type level but negatively correlated at the individual level. The impact of this different correlation structure is apparent in the distribution of the fraction of positive associations obtained in the analysis of randomly permuted expression datasets (Figures 3A and 3B). Random distributions were matched with observed data by the number of significant associations per neighbour. Compared with these random distributions, neighbours enriched in BCT DA have more positive associations and fewer negative or mixed patterns than expected. Neighbours enriched in WCT DA have a sign distribution that is similar to the random expectation. The distribution of the fraction of positive associations was also performed for drivers enriched in BCT and WCT NA (Figures 3C and 3D). For both types of analysis, drivers enriched in NA tend to have a mixed pattern of associations. Drivers enriched in BCT NA have a shift towards higher fractions of positive associations when compared with the random expectation (Figure 3C). This is in agreement with the predominance of neighbours enriched in positive BCT DA. Drivers enriched in WCT NA tend to have a mixed signal pattern, with slightly more negative than positive associations (Figure 3D). Drivers enriched in WCT NA with coherently positive or negative associations are less frequent than randomly expected.

### Neighbours enriched with driver associations tend to have cancer survival impacts that are coherent with the association signs

In order to verify whether associations between neighbour expression and driver mutation status had a significant impact on the clinical outcome, we assessed the impact of neighbour expression on overall survival across 21 cancer types according to a Human Pathology Atlas resource (26, 27). First, we observed that neighbours enriched BCT and WCT DA had a significantly higher proportion of neighbours with at least two prognostic survival associations (SA) in two different cancer types (Table 1). Next, we found that, in both BCT and WCT analyses, neighbours with an unfavourable prognostic SA in more cancer types tended to have more positive driver associations (Figure 4A). This is a sign-coherent trend, as positive neighbour-driver associations indicate that high neighbour expression promotes the tumorigenic effects of driver mutations and, consequently, poorer survival outcomes. Conversely, neighbours with more favourable prognostic SA also tended to have more negative BCT DA (Figure 4B). However, a similar trend was not observed for negative WCT DA. Finally, we classified neighbours into four classes according to their SA: 1) neighbours that do not have any SA, 2) neighbours that only have favourable SA, 3) neighbours that only have unfavourable SA and 4) neighbours that have both favourable and unfavourable SA. We then tested if these four classes were more or less frequently found in sets of neighbours enriched in positive, negative or mixed DA, both for BCT (Figure 4C) and WCT analyses. Neighbours enriched in positive BCT DA were significantly enriched in unfavourable SA, and showed lower than expected proportions of neighbours with favourable or no SA (Figure 4C). Coherently, neighbours enriched in negative BCT DA were significantly enriched in favourable SA and had lower than expected proportions of neighbours with unfavourable or no SA (Figure 4C). For the WCT type of associations, only neighbours enriched in mixed WCT DA were enriched in neighbours with both favourable and unfavourable SA (Figure 4D). Globally, these results validate the impact of driver-neighbour associations in disease outcome. Although neighbours with mixed WCT DA can have a relevant role in conditioning the effect of several driver mutations, their context dependent effects turn them less attractive as candidate drug targets. Therefore, we selected two lists of neighbours for further exploration as potential drug targets: 1) 119 neighbours enriched in positive BCT DA with at least two unfavourable SA (Supplementary file 2) and 2) 25 neighbours enriched in negative BCT DA with at least two favourable SA (Supplementary file 3).

**Figure 4.**
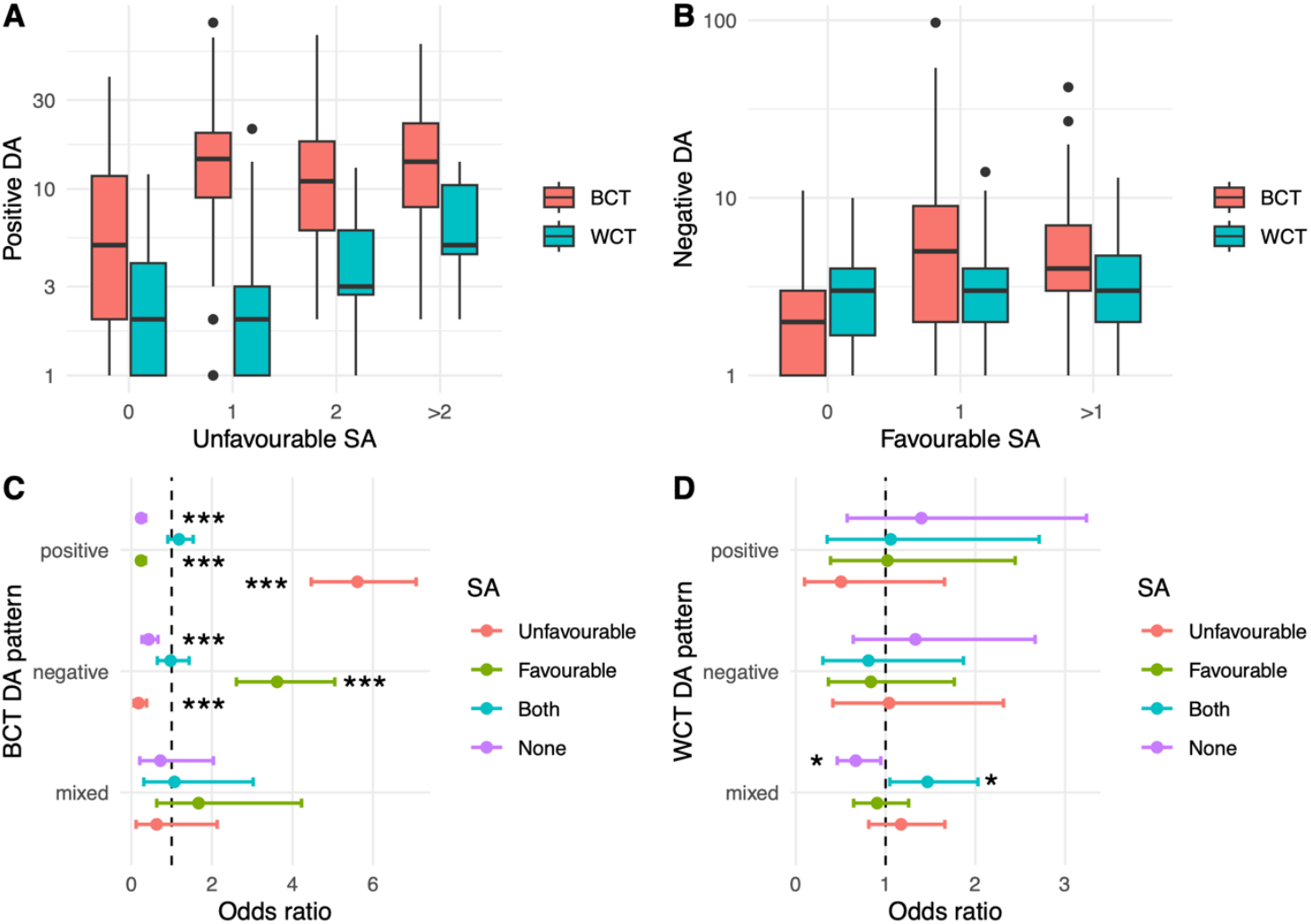
Association between driver and survival associations. (A) Boxplots representing the distribution of significant driver associations (DA) with a positive sign as a function of the number of cancer types for which a significant unfavourable survival association (SA) was detected, for both between cancer types (BCT) and within cancer types (WCT) analyses. (B) Boxplots representing the distribution of significant DA with a negative sign as a function of the number of cancer types for which a significant favourable SA was detected, for both BCT and WCT analyses. (C) Odds ratios for the association between neigbours with a particular SA pattern (only unfavourable SA, only favourable SA, both favourable and unfavourable SA and no SA) and neighbours enriched in BCT DA with a particular sign pattern (positive – 80% or more of positive DA, negative – 20% or less of negative DA, mixed – between 20% and 80% of positive DA). (D) Odds ratios for the association between neigbours with a particular SA pattern (only unfavourable SA, only favourable SA, both favourable and unfavourable SA and no SA) and neighbours enriched in WCT DA with a particular sign pattern (positive – 80% or more of positive DA, negative – 20% or less of negative DA, mixed – between 20% and 80% of positive DA). In (C) and (D), error bars represent 95% confidence intervals for the odds ratios. ^*^ represents p<0.05 and ^***^ represents p<0.0001 for hypergeometric tests (Null hypothesis: Odds ratios are equal to 1). Odd ratios significantly greater than 1 mean that the proportion of neighbours with a particular SA pattern within neighbours enriched in DA with a particular sign pattern is greater than expected by chance.

### Most neighbours simultaneously enriched in driver and survival associations have evidence of causing effects in cancer cells that are coherent with driver association sign

The candidate neighbour sets were selected through significant statistical associations with driver mutation status and overall survival. Both associations were evaluated with observational data from a large cancer patient cohort. Therefore, these associations alone are not enough to prove that these neighbours have any causal effect on cancer cells. A fraction of these neighbours are themselves cancer drivers (35 out of 119 positively associated neighbours and 6 out of 25 negatively associated neighbours). Being cancer drivers implies that changes in their activities induce causal effects on cancer cells. Surprisingly, three tumour suppressor genes (*BUB1B, FANCA* and *POLD1*) are among the neighbours enriched in positive associations with drivers and decreased survival in multiple cancer types. The pro-tumorigenic effect of high expression of some of these genes has been previously reported (29, 30). These genes are involved in DNA repair pathways. Loss of function mutations in DNA repair genes may lead to accumulation of mutations and chromosomal instability, promoting cancer development and explaining their tumour suppressor roles. But when they are not mutated, higher expression may be associated with more aggressive tumours, where these pathways may be activated to prevent excessive chromosomal instability that could lead to cell death (31). Higher DNA repair capabilities may also increase the resistance to some therapies, which may cause decreased survival times (32).

For neighbours not previously identified as cancer drivers, we conducted a bibliographic search to find experimental evidence (functional tests using overexpression, knockdown, or knockout in cancer cell lines) of causal effects that were coherent with the dominant sign of driver and survival associations. We successfully identified such experimental evidence for 88% (74 out of 84, Supplementary file 2) of positively associated neighbours and for 37% (7 out of 19, Supplementary file 3) of negatively associated neighbours.

We note that, for some neighbours, we were able to identify evidence with opposing effects. As an example, *PTGR2*, a neighbour enriched in negative driver associations, has been shown to promote gastric cancer (33) while having favourable SA in lung and renal cancer. These dual roles could be justified by a mixture of positive and negative driver associations. This is not the case for *PTGR2*, for which only negative driver associations were detected. This suggests that *PTGR2* may have other PPIs with cancer-related proteins that explain its pro-tumorigenic role in gastric cancer.

Finally, we searched the Open Target platform (34) and found that 28 of our selected neighbours (25 neighbours enriched in positive driver and survival associations and 3 enriched in negative driver and survival associations, Supplementary files 2 and 3) are known drug targets. 19 out of these 28 were not previously recognized cancer drivers. 18 are targeted by drugs developed for cancer treatment, confirming in this way the therapeutic potential of neighbours enriched in driver associations. Interestingly, 10 neighbours (*CX3CL1, E2F1, LAMA3, MIF, RPL28, RPL35, RPL7A, RPLP0, RPS15A, RPS21*) were targeted only by drugs for non-cancer related diseases, suggesting an opportunity for drug repurposing.

## Discussion

This study investigated the relationship between driver mutations, neighbour gene expression, and patient survival in cancer. We were able to identify significant associations between neighbour expression and driver mutation status that were not explained by a direct mutation effect. These associations were detectable both in comparisons between and within cancer types. Moreover, we could identify neighbours that had more driver associations than expected by chance. Many of these enriched neighbours tended to have sign-coherent associations with driver mutation status, either mostly positive or mostly negative associations. One key finding was the significant correlation between the number of driver associations and the number of sign-coherent survival associations for each neighbour. This correlation is particularly strong for neighbours enriched in BCT driver associations. Neighbours enriched in driver associations were significantly more likely to have multiple survival associations, particularly for neighbours with positive associations, where high neighbour expression correlated with both increased driver mutations and poorer survival.

The fact that direct neighbours of cancer drivers are associated with patient overall survival is not surprising, as it has been observed that physical interactors of cancer drivers have an increased probability of being cancer drivers themselves (13). There are also previously known examples of neighbours that condition the effects of driver mutations, such as EGFR and c-CBL (35) or TP53 and MDM2 (36). In both examples, the neighbour proteins are E3 ubiquitin ligases that control the degradation rate of the driver proteins, therefore regulating their abundance and activity. The novelty in this work resides in the finding that such neighbour effects can leave a statistical signature in genomic and transcriptomic datasets of cancer patient cohorts. This signature allows for the systematic discovery of neighbours with significant driver associations that can be relevant prognostic biomarkers or therapeutic targets.

Not all neighbours that condition driver mutations effects are discoverable in this way. For the statistical driver association to be detectable, it is necessary that the neighbour has a natural expression variability, either between tissues or between individuals. On the other hand, similar statistical associations with driver mutation status and with patient overall survival can also exist for other proteins that are not neighbours of cancer drivers. The advantage of detecting an association for a neighbour protein is that the PPI supports the existence of a molecular mechanism explaining the statistical association.

Our study identified 119 neighbours enriched in positive driver associations with multiple unfavourable survival associations and 25 neighbours enriched in negative driver associations with multiple favourable survival associations. These neighbours represent promising candidates for further investigation as potential drug targets. In fact, 18 of them are already known drug targets for cancer therapy. Additionally, a literature search revealed experimental evidence supporting the causal roles of a substantial proportion of these neighbours.

We opted to focus on neighbours that have most of their driver associations with the same sign. Their role in cancer development is less context-dependent, facilitating the detection of significant survival associations. Neighbours with a mixture of both positive and negative driver associations are not less valid. In fact, they can be fundamental to fully understand the role of drivers that behave both as tumour suppressors or as oncogenes, depending on the cellular context. But to untangle these context-dependent effects, either more data or experimental approaches are needed.

While the statistical associations observed in this study provide compelling evidence for the role of proteins that interact directly with cancer driver proteins, further experimental validation is essential to establish causality. Future research should focus on characterising the functional roles of the identified neighbours and exploring their potential as therapeutic targets. This could involve in vitro and in vivo studies to assess the impact of modulating neighbour gene expression on cancer cell growth, survival, and response to therapy. The impact of neighbour modulation should be stronger when the cancer cells harbour mutations in drivers interacting with that neighbour. The confirmation of these neighbours as drug targets, especially effective for subsets of patients carrying particular driver mutations, would expand the toolkit of targeted therapies.

It is important to acknowledge certain limitations of this study. The analysis relied on observational data from the TCGA pan-cancer cohort. The main advantage of using observational data is that measurements are made in real patient tissues instead of animal or cell models. However, it is not possible to control all confounding variables in observational studies. This limitation implies that we cannot control for a situation in which some individuals were exposed to unknown factors that could affect simultaneously driver mutations and neighbour expression and, therefore, generate an artificial association between driver mutation and neighbour expression.

Using observational data also implies that the number of cases with a particular combination of attributes is not controlled by the researchers. This limits the application of more complex statistical models. In the linear models used to detect associations between driver mutation status and neighbour expression, it would be interesting to include interaction terms between mutation status and cancer type. This would allow us to detect more driver-neighbour associations that would be conditional on the cancer type. But given that for most drivers there is only a reduced number of mutated individuals among the less prevalent cancer types, the statistical power is not sufficient for a precise estimation of the referred interaction coefficients.

RNA-seq data was used to estimate the abundance of neighbour proteins in cancer cells. Correlation between mRNA and protein levels can be rather low, especially when comparing values of different genes in the same tissue or cell type (37). However, when comparing values for the same gene across different samples, mRNA-protein correlations are much higher (38). In the present study, mRNA expression values being compared are always for the same neighbour gene across different individuals and cancer types. Therefore, we expect that most of these changes in gene expression reflect similar changes in protein levels.

Other limitations arise due to tumour heterogeneity and the diverse composition of the tumour microenvironment. RNA-seq was performed on the bulk of cells collected from the tumour tissue. We cannot be sure that all tumour cells carry the same mutations and have similar gene expression profiles. And part of the expression signal may come from non-tumour cells that are infiltrated in the tumour tissue. This issue could be solved by utilizing single-cell genomic sequencing and mRNA sequencing for each tumour sample. In this way, we could identify neighbour gene expression in the same cells that carried a mutation in the interacting driver. Besides the additional layer of complexity in the data analysis, we are not aware of the availability of this type of data for large cohorts of cancer patients.

The PPI datasets used might have their own limitations. It is possible that interaction data is incomplete, meaning that drivers may have unknown interactions with other proteins. Additionally, some interactions are only established with mutated drivers (39) and are not included in the interaction databases used here. Although our focus was on protein physical interactions, relevant interactions may be established with other types of molecules, such as non-coding RNAs. Finally, we only considered binary interactions. Some context-dependent effects of driver mutations may be explained only by higher-order interactions, involving more than one interactor simultaneously.

Despite these limitations, the findings of this study provide valuable insights into the complex interplay between drivers and their neighbours. It also demonstrates a new approach to analyse genomic and transcriptomic data from cancer patient cohorts that uncovers neighbours enriched in driver associations as potential therapeutic targets and prognostic biomarkers.

## Materials and Methods

### Cancer drivers

A list of canonical and candidate cancer driver genes was retrieved from the Network of Cancer Genes & Healthy Drivers 7.1 (19). This list was used to search for protein physical interactors and to detect driver-neighbour associations (meaning that neighbour gene expression was statistically associated with driver mutation status). To evaluate if neighbours enriched in driver associations had any known cancer driver role, we retrieved the lists of cancer driver genes available in COSMIC Canger Gene Census (40), Intogen (2) and OncoKB (41, 42). Additionally, for the final lists of neighbours enriched in driver associations and with multiple survival associations, we searched the Disgenet database (43) for evidence of association with neoplastic diseases with association type “CausalMutation”.

### Protein physical interaction data

Human PPI were retrieved from five sources: String (only PPI with a confidence score equal or greater than 0.4) (21), Biogrid (20), HuRI (22), Omnipath (23) and APID (24). Interactions involving complexes were expanded to include all possible pairwise interactions. Protein physical interactors (neighbours) of each cancer driver were extracted from all five interactome sources. General characteristics if these networks are shown in Supplementary table 1.

### Cancer patient data

Cancer patient data from the TCGA Pan-Cancer cohort was accessed through the UCSC Xena online tool (https://pancanatlas.xenahubs.net) (44). In particular, gene-level non-silent somatic mutation data (18), batch effects normalised mRNA data, sample phenotype data and patient clinical data (45) were retrieved. Patients with more than 72 mutated cancer driver genes were discarded as they were considered mild outliers using the Tukey’s fence method (46). Missing values in the gene expression data were imputed with the minimum expression value observed for the respective gene across all patients.

### Tumour mutational burden

Tumour mutational burden (TMB) is a measure of the total number of somatic mutations found in a tumour specimen. It is commonly used as a biomarker useful to predict the outcome of certain immunotherapies. TMB can be characterized experimentally using different methodologies that look for nonsynonymous mutations in coding regions and oncogenic drivers, including whole exome sequencing (WES) (47). In our work, we calculated the TMB for each tumour specimen (patient) using a dataset which includes all somatic mutation calls (SNPs and INDELs). First, this dataset was filtered to remove silent mutations—we considered as silent mutations mutations in noncoding regions—and then we counted all nonsynonymous mutations for each sample. For the BCT analysis we computed the mean mutational burden per cancer type. TMB values, both for individual patients and cancer type averages, were converted with a *log*_10_ transformation.

### Comparison of paired tumour and normal samples

596 patients in the pan-cancer cohort had paired tumour and normal tissue samples. For each of these patients, we computed the difference in gene expression between the tumour sample and the matching normal tissue. For each driver-neighbour PPI, we fitted a linear regression model with fixed effects (Equation 1).

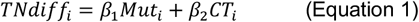

The dependent variable was the tumour-normal neighbour gene expression difference (*TNdiff*_*i*_). The independent variables were 1) the mutation status of the driver (*Mut*_*i*_) and 2) the cancer type (*CT*_*i*_) of each patient *i*. Only data from cancer types that had at least one patient with a positive mutation status in at least 2 cancer types were included in the model. A significant (p<0.05) regression coefficient of the driver mutation status (*β*_*i*_) was considered as evidence that the driver mutation induced a change in the expression of the neighbour. These driver-neighbour PPIs were excluded from further analyses.

### Pathway analysis

Driver mutations might have an impact on neighbour gene expression if the driver protein directly regulates signalling pathways and transcription factors (TF) that target the neighbour gene. To exclude this type of association, we built a regulatory network by combining two directed networks: a network of signalling pathways from Omnipath (23) with a TF-target gene network from DoRothEA (25) (Supplementary table S1). For each driver-neighbour physical PPI, we computed the shortest path distance between the driver and the neighbour so that the final interaction was between a TF and the neighbour gene. A distance of 1 means the driver is a direct TF of the neighbour. Driver-neighbour pairs with a shortest distance of 1 or 2 were more likely to have a significant difference in the tumour/normal tissue comparison, and thus were excluded from the analysis.

### Statistical associations between neighbour expression and driver mutation status

We used data from 7317 patients (which passed the driver mutation outlier filter and had mutation and gene expression data for tumour samples) to compute, for each driver-neighbour PPI, the Spearman correlation coefficient between 1) the average neighbour expression per cancer type and 2) the fraction of patients with mutated driver divided by the average TMB per cancer type. In this way we can avoid detecting false correlations where the higher driver mutation frequency in some cancer types is just due to a higher TMB of the individuals with those cancer types. A statistical test was applied to each driver-neighbour PPI to evaluate if the computed coefficient was significantly different from 0. Previous to the analysis, we conducted a power analysis with simulated data and concluded that to achieve a statistical power of 0.7, the correlations should only be computed when there are at least 10 cancer types with a driver mutation frequency greater than 0. Driver-neighbour PPIs with a significant correlation (p<0.05) were considered statistically associated in a between cancer type comparison (BCT).

Using the same data, we fitted, for each driver-neighbour PPI, a linear regression model with fixed effects (Equation 2).

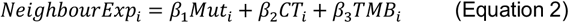

The dependent variable was the neighbour gene expression (*NeighbourExp*_*i*_) and the independent variables were 1) the driver mutation status (*Mut*_*i*_), 2) the cancer type (*CT*_*i*_) and 3) the tumour mutation burden (*TMB*_*i*_) for each patient *i*. Previous to the analysis, we conducted a power analysis with simulated data and concluded that to achieve a statistical power of 0.7, the linear model should only be fitted when there are at least 3 cancer types with at least one patient with a driver mutation. A significant (p<0.05) regression coefficient of the driver mutation status (*β*_*i*_) was considered as evidence that the driver-neighbour PPI was statistically associated in a comparison within cancer types (WCT).

### Significant neighbour identification through random permutations

To evaluate the statistical significance of the number of driver-neighbour associations per neighbour protein, the computation of BCT and WCT associations was repeated 100 times with randomly permuted gene expression datasets. Gene expression values were organized in a matrix where each row corresponded to an individual and each column corresponded to a gene. Randomly permuted matrices were produced by randomly shuffling the row order without changing the column orders. In each shuffle, the number of significant driver associations (*x*_*i*_) of each type (BCT or WCT) for each neighbour *i* was recorded, allowing the computation of the average (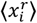) and standard deviation (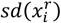) of this number according to a random null model. These parameters were used to compute z-scores (Equation 3) for the original observations and for each of the 100 analyses with randomly permuted data.

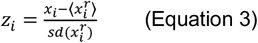

For any given z-score threshold (*t*) we computed the False Discovery Rate (FDR, Equation 4) using the average number of neighbours with a z-score greater than the threshold in the 100 analyses with the randomly permuted data (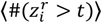) as an estimate of the expected false positives.

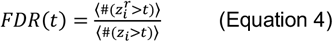

Equation 4 estimates FDR by excess, as the estimated number of false positives is done with the total number of neighbours instead of using the unknown proportion of neighbours that follows the null hypothesis (*H*_0_: the neighbour does not have more driver associations than expected by chance). For each type of driver association (BCT or WCT), we determined the lowest value of *t* for which *FDR*(*t*) < 0.1.

### Survival analysis

Cancer prognostic data was downloaded from the Human Protein Atlas, containing Log-rank p values for Kaplan-Meier analysis of correlation between mRNA expression level and patient survival in prognostic cohorts based on 21 cancers from TCGA and validation cohorts for 10 of the cancers. Details of the analysis are described in previous publications (26, 27). According to the authors, genes were designated as prognostic genes if they had log-rank p values less than 0.001. Benjamini and Hochberg procedure (48) was applied to the 219519 p-values available in the dataset. The p-value threshold of 0.001 corresponded to an FDR of 0.013. 22640 gene-cancer type prognostic associations were considered using this threshold.

### Tissue specificity

We used the Tau Tissue Specificity Index (49) to characterize the tissue specificity of genes across different tissues. The tau index (*τ*), is defined as (Equation 5)

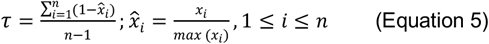

Where *n* is the number of tissues and *x*_*i*_ is the expression of the gene in tissue *i*. We applied the tau index to the Human Protein Atlas (HPA) Consensus RNA dataset. This dataset summarizes HPA (50) and GTEx (51) transcriptomics data across 50 tissues using a normalized TPM value (nTPM). To eliminate similar tissues that could impact the tissue specificity index calculation (e.g. the brain is represented with 8 different regions), we processed the dataset into 41 distinct organs by taking the median nTPM of tissues in the same organ. Finally, nTPM values were *log*_2_ transformed, and an expression cutoff was applied: all genes with expression below 1 nTPM were removed.

## Code availability

All the code used to perform the analyses presented in this manuscript is available at https://github.com/GamaPintoLab/driver_neighbours. All the data used in this work was obtained from public sources. For reproducibility purposes, raw data files used are available at https://doi.org/10.5281/zenodo.14284407.

## Acknowledgments

Work supported by UID/04046/2025 – Instituto de Biosistemas & Ciências Integrativas Centre grant from FCT, Portugal.

