## Supplementary tables and figures for "When neighbours play a role: a systems-level analysis of protein interactions conditioning cancer driver mutation effects"

Table S1 – General properties of networks used in this work.

| Network | Nodes | Edges | Drivers | Neighbours | Average degree | Average degree of drivers | Average degree of nodes |
|---|---|---|---|---|---|---|---|
| APID (PPI) | 16994 | 262291 | 2908 | 14390 | 30.87 | 41.94 | 33.12 |
| BIOGRID (PPI) | 19694 | 868997 | 3053 | 15300 | 88.25 | 138.08 | 102.4 |
| Dorothea (TF-target) | 11704 | 47306 | 2321 | 10158 | 8.08 | 17.75 | 8.90 |
| HURI (PPI) | 9050 | 63584 | 1305 | 7688 | 14.05 | 13.64 | 14.55 |
| OMNIPATH (Signalling) | 10408 | 106749 | 2256 | 9503 | 20.51 | 32.86 | 21.63 |
| STRING (PPI) (conf ⩾ 0.4) | 15632 | 410424 | 2789 | 13493 | 52.51 | 74.95 | 54.10 |
| Driver-Neighbour fused network (PPI) | 15683 | 455604 | 3081 | 15464 | 58.10 | 147.88 | 29.46 |

Table S2 – Comparison between paired analysis (Tumour—Normal tissue comparison) and signalling / transcriptional regulatory network analysis.

| Shortest path length between Driver and Neighbour in signalling / transcriptional regulatory network | Paired analysis | | Fraction of significant Driver—Neighbour pairs | P-value (hypergeometric test) |
|---|---|---|---|---|
| | Not significant | Significant | | |
| 1 | 977 | 157 | 0.138 | 5.30e-21 |
| 2 | 19747 | 1540 | 0.072 | 8.63e-11 |
| 3 | 54702 | 3657 | 0.063 | 0.353 |
| 4 | 34986 | 2162 | 0.058 | 0.999 |
| 5 | 8346 | 529 | 0.060 | 0.862 |
| 6 | 1626 | 91 | 0.053 | 0.944 |
| 7 | 263 | 11 | 0.040 | 0.925 |
| 8 | 14 | 1 | 0.067 | 0.240 |

Table S3 – Comparison between BCT associations with all cancer types (All) and BCT associations assessed by splitting cancer types (CTs) into high and low driver mutation frequency groups.

| Driver-Neighbour pairs | Comparison | Fraction with concordant sign | Correlation coefficient |
|---|---|---|---|
| Significant BCT association | All vs CTs with low mutation frequency | 0.831 | 0.665 |
| | All vs CTs with high mutation frequency | 0.833 | 0.687 |
| Non-significant BCT association | All vs CTs with low mutation frequency | 0.608 | 0.336 |
| | All vs CTs with high mutation frequency | 0.623 | 0.373 |

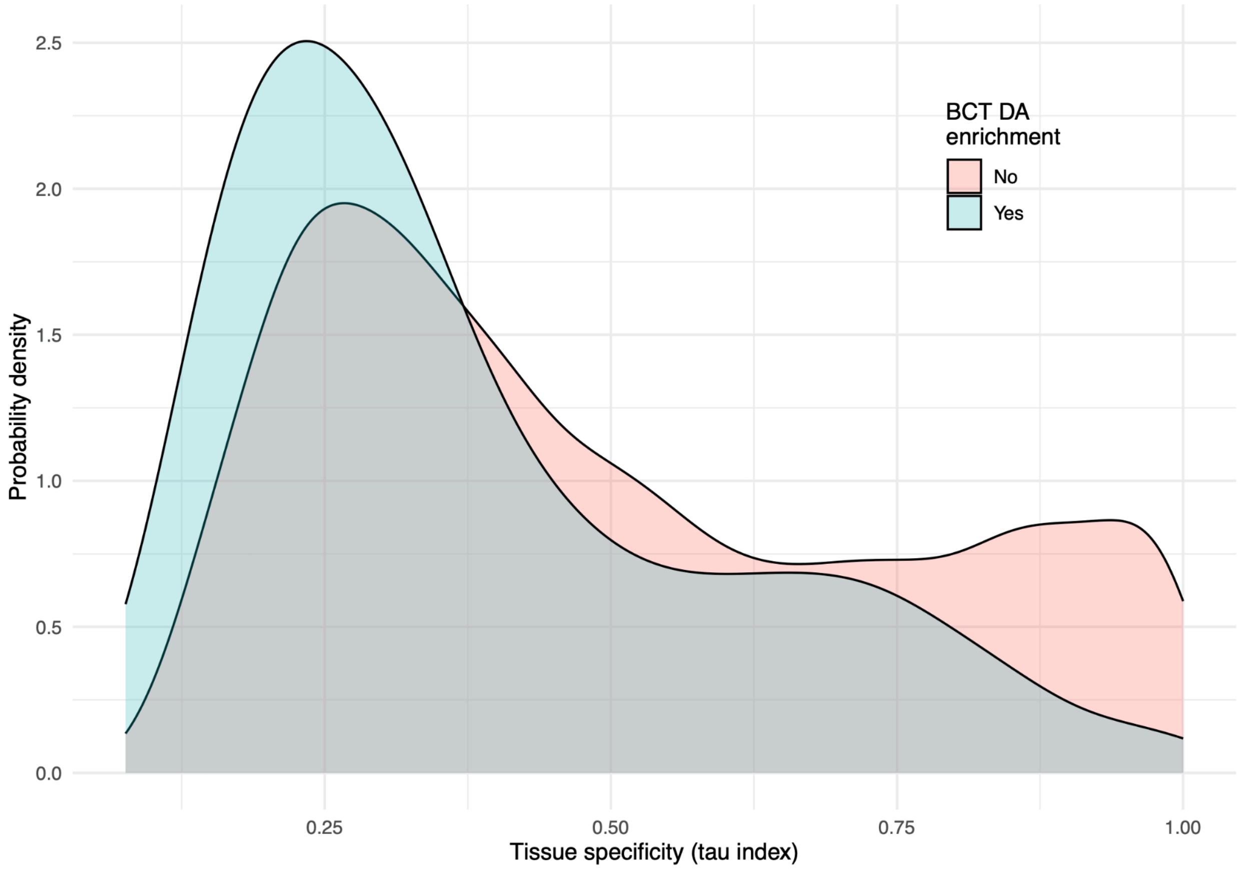


Figure S1 – Tissue specificity of neighbours with between cancer types (BCT) driver associations (DA). Probability density distribution of the tau index for neighbours enriched (or not) in BCT DA. The tau index measures the tissue specificity of gene expression. Tau values close to 1 means that the gene is expressed in only 1 or a few tissues. Tau values close to 0 mean that the gene is uniformly expressed across all tissues. Tau indexes were computed with tissue gene expression data publicly available from the Human Protein Atlas resource.

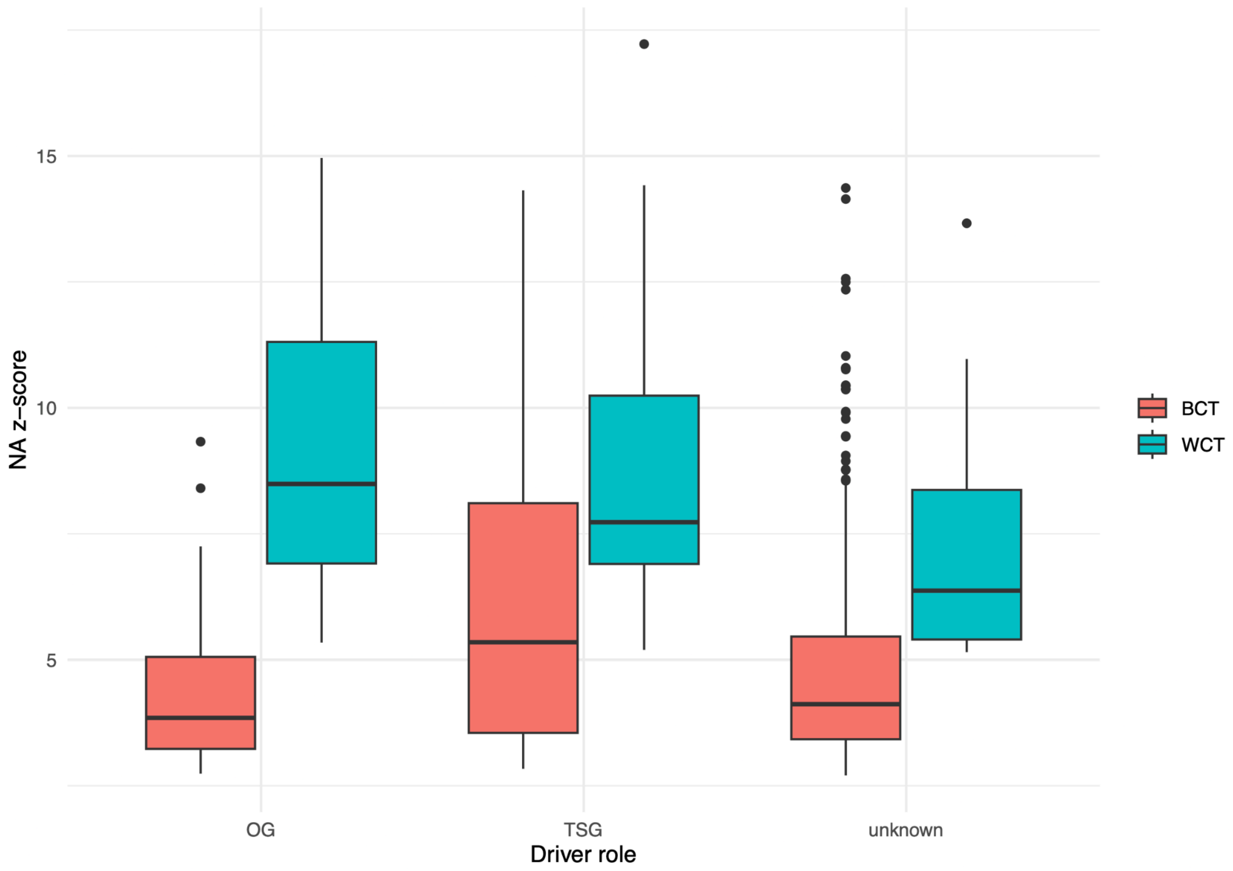


Figure S2 – Enrichment in neighbour association (NA) for drivers with known roles as Oncogenes or Tumour suppressor genes. Boxplots of the NA z-score for drivers enriched in between cancer types (BCT) NA or within cancer types (WCT) NA. NA z-score is the difference between the number of significant neighbour associations a driver has and the randomly expected number of significant associations in units of the standard deviation of the random distribution of the number of significant associations. Higher z-scores mean that the driver has more significant neighbour associations than were expected by chance.
